# Beta-2-microglobulin stimulates neutrophil phagocytosis of bacteria and apoptotic cells

**DOI:** 10.1101/2025.10.30.685267

**Authors:** Sofie Espersen Poulsen, Michal Magda, Anna Blom, Mogens Holst Nissen

## Abstract

Efficient clearance of dead cells and pathogens is essential for survival and requires both the innate and adaptive immune systems. Polymorphonuclear leukocytes (PMNs, predominantly consisting of neutrophil granulocytes) constitute an important first line of defense. These highly specialized cells can phagocytose pathogens and clear apoptotic cells through efferocytosis.

Beta-2-microglobulin (β2m) serves as the light chain of major histocompatibility complex class I (MHC I) molecules, associating non-covalently with the heavy transmembrane chain that binds and presents antigenic peptides to CD8^+^ T cells. This represents the canonical role of β2m. β2m is also found in the granules of PMNs and is released into the extracellular space during degranulation. However, a specific function for β2m in the context of PMN function or degranulation has not yet been identified.

We now present evidence that β2m is of importance for both phagocytosis of pathogens and efferocytosis of dead cells by PMNs. The addition of exogenous β2m (50 mg/l) to PMNs in the presence of latex beads increased the phagocytic activity from 23% to 31%. Furthermore, both β2m and desLys58-β2m (dK58β2m) enhanced phagocytosis of Gram-negative and -positive bacteria by more than 3.6-fold, though no effect was observed with zymosan bioparticles.

Maintaining tissue homeostasis requires the continuous generation of new cells and the efficient clearance of apoptotic cells through efferocytosis. Treatment with β2m or dK58β2m led to a dose-dependent increase in efferocytosis of apoptotic Jurkat cells, reaching up to a two-fold enhancement. This effect was comparable to that obtained by GM-CSF, used as a positive control. In all cases, cytochalasin D blocked β2m-mediated uptake in PMNs.

These data demonstrate that β2m can be of importance in phagocytosis of bacterial pathogens as part of the innate immune response and tissue homeostasis by removing dead cells by efferocytosis.

## Introduction

Beta-2-microglobulin (β2m) constitutes the invariant light chain of the tri-molecular major histocompatibility complex class I (MHC I), where it non-covalently associates with the transmembrane heavy chain responsible for presenting peptide antigens to CD8^+^T cells^1,2^. In its soluble form, β2m is present in various biological fluids^3^, and is also stored within neutrophil granules and platelets^4–7^. Additionally, it can be secreted into the extracellular environment by activated immune and epithelial cells^8,9^. Elevated serum β2m levels have been reported in patients with a range of inflammatory^10–13^, malignant^14–16^, viral^17,18^, and renal diseases^19^. β2m is an important prognostic factor for several diseases, and high serum levels often correlate with poor prognosis^20–23^.

A modified variant of β2m, later characterized as desLys58-β2m (dK58β2m), was first found in the serum of patients with systemic lupus erythematosus and rheumatoid arthritis^24^. Subsequent studies revealed that dK58β2m is generated via a two-step proteolytic process: native β2m is first cleaved at the C-terminal side of lysine 58 by the complement component C1s, followed by removal of lysine 58 by a carboxypeptidase B-like activity^25,26^. This cleavage induces a conformational change in β2m, conferring amyloidogenic properties that increase its propensity to aggregate and form fibrils^27–29^. The presence of dK58β2m has also been reported in various cancer types^30–33^. Only a limited number of studies have investigated the biological significance of the cleavage. This includes augmentation of specific cytotoxic activity in a one-way mixed lymphocyte culture^34^, induction of nitric oxide generation and apoptosis in combination with interferon-γ (IFN-γ) in myeloid cell lines^35^. Notably, a recent study demonstrated that dK58β2m exacerbates mitochondrial dysfunction, mtROS production, and activation of the cGAS-STING pathway in macrophage-like cells, particularly in the presence of IFN-γ^36^.

Neutrophil granulocytes are the most abundant leucocytes in human blood, constituting about 50-70% of all circulating immune cells. As primary responders of the innate immune system, they are critical for the elimination of extracellular pathogens, particularly via phagocytosis, a process involving recognition, internalization, and destruction of microbial targets^37^. Following pathogen clearance, neutrophils undergo apoptosis and are subsequently cleared via efferocytosis by phagocytic cells such as macrophages, a process essential for resolving inflammation and preventing tissue damage^38,39^. Although traditionally seen as passive substrates for efferocytosis, recent evidence suggests that neutrophils may actively influence or participate in their own clearance^40,41^.

β2m localized within neutrophil granules is released into the extracellular environment during degranulation, yet the functional role of this intragranular pool of β2m remains unclear. Other granule proteins, such as Cathepsin G, have been shown to regulate neutrophil effector functions through autocrine or paracrine mechanisms^42^. Given this, we aimed to investigate whether β2m or its cleaved form, dK58β2m, influence neutrophil-mediated phagocytosis and efferocytosis.

In this study, we demonstrate that both β2m and dK58β2m exert stimulatory effects on phagocytosis and efferocytosis by PMNs. β2m significantly enhanced the uptake of latex beads as well as both Gram-positive *Streptococcus pyogenes* AP1 and Gram-negative *Acinetobacter baumannii* KR792, whereas dK58β2m only increased phagocytosis of bacterial targets but not of latex beads. Furthermore, both proteins stimulated the efferocytic uptake of apoptotic Jurkat cells by PMNs, highlighting a potential modulatory role of β2m and its cleavage product in neutrophil functions. This points towards an involvement of β2m in the early innate immune response and in the resolution of inflammation.

## Materials and methods

### Protein expression, cleavage, and purification

Human recombinant β2m was produced in the ExpiCHO™ expression system (Thermo Fisher - Scientific, Gibco™, #A29133). The dK58β2m variant was created by cleavage of β2m, and all protein variants were purified as described previously with minor modifications^36,43^. Initially, both proteins were separated on a G75 Sephadex gel filtration column. The dK58β2m variant was subsequently purified by ion exchange chromatography on a Resource Q column. Finally, the protein samples were up-concentrated by ammonium sulphate precipitation and subsequently passed through PD-10 desalting columns.

### Cells

The human T lymphoblast cell line Jurkat (TIB-152™, ATCC®) was cultured in RPMI 1640 medium supplemented with 1% L-glutamine, 1% penicillin and streptomycin, and 10% fetal bovine serum (FBS; Corning, #35–079-CV) at 37°C with 5% CO_2_. For efferocytosis assays, Jurkat cells were treated with 0.8 µM staurosporine (Sigma-Aldrich, #S5921) for 24 h to induce apoptosis. The degree of apoptosis was assessed using annexin V-FITC (Nordic Biosite, #640906) and 7AAD (Nordic Biosite, #420403) staining according to the manufacturer’s recommendations. On average, 7±6% were double negative (e.g. live), 30±13% were annexin V positives (e.g. early apoptotic), 62±16% were double positives (e.g. late apoptotic) and 0.7±0.8% were 7AAD positives (e.g. necrotic or another type of cell death) (Supplementary figure 1)

### Bacteria

*Streptococcus pyogenes (S. pyogenes)* strain AP1 (strain 40/58 serotype M1)^44^ was grown in Todd-Hewitt broth medium (Oxoid, #CM0189) at 37°C with 5% CO_2_, while *Acinetobacter baumannii (A. baumannii)* clinical isolate KR792^45^ was grown on a blood agar plate at 37°C. Following overnight incubation, AP1 was diluted in fresh medium to OD_600_=0.1 and grown for approximately 2 h until it reached an exponential growth phase at OD_600_=0.3-0.4. Simultaneously, an overnight KR792 was subcultured into a fresh blood agar plate and incubated for an additional 3 h. After incubation, AP1 and KR792 were collected, washed with PBS, and diluted to the required concentration. The bacteria were kept on ice until used in the assay.

### Normal human serum

Normal human serum (NHS) was obtained from the blood of 14 healthy donors and stored at −80°C as described previously^46^. Collection of serum was approved by the Swedish Ethical Review Authority (permit number 2023-05543).

### Isolation of human peripheral blood PMNs for phagocytosis assays (using latex beads or zymosan) and for efferocytosis assays

Human PMNs were isolated from heparinized whole-blood samples collected from healthy donors attending the Copenhagen University Hospital blood bank, Denmark. In accordance with Danish legislation and ethical guidelines, donor identities remained completely anonymous, with samples only labeled by age and gender. Isolation of PMNs was performed as described previously^47^. In short, whole blood was mixed with dextran T-500 (Sigma-Aldrich, #31392) at a 5:1 ratio, allowing erythrocytes to sediment for 1 h at room temperature. The leucocyte-enriched plasma was then carefully layered over a Lymphoprep™ gradient (Nordic Biosite, #318-1856) in a 1:1 ratio and spun down. The resulting pellet, containing PMNs and residual erythrocytes, was collected, and erythrocytes were lysed by exposure to hypotonic saline (0.2%) for 1 minute, followed by the addition of 1.6% saline to restore isotonic conditions. Finally, the PMNs were resuspended in HBSS, and live cells were counted. The cells were used for experiments immediately after isolation.

### Isolation of PMNs for phagocytosis assays with bacteria

Human neutrophils were isolated following a previously described method^48^. Blood from healthy donors was collected under ethical approval (permit number 2023-05543) granted by the Swedish Ethical Review Authority. Briefly, PMNs were separated using Histopaque 1119 (Sigma-Aldrich, #11191). The fraction containing PMNs was collected, washed with PBS, and further separated using a gradient of Percoll (GE Healthcare, #1708-0891-01) and RPMI 1640 (Cytiva, #SH30027.01) supplemented with 10 mM HEPES (Cytiva, #SH30237.01). After centrifugation, the distinct layer containing PMNs was collected and washed with PBS containing 0.5% BSA (Saveen Werner, #B2000-100). PMNs were then suspended in RPMI-HEPES. Live cells were counted, diluted to 1×10^6^/ml, and kept at RT until used in the assay.

### Flow cytometric analysis of phagocytosis of latex beads and zymosan bioparticles

A total of 2.5×10^5^ PMNs were incubated with 1 µm Red FluoSpheres™ Carboxylate-Modified Polystyrene Microspheres (Latex beads; Thermo Fisher Scientific, #F8821) either alone or in combination with 50 µg/ml β2m or dK58β2m for 4 h at 37°C, 5% CO_2_. Latex beads were added at a PMN-to-particle ratio of 1:10. PMNs pre-treated with 10 µg/ml cytochalasin D (Thermo Fisher Scientific, #PHZ1063) for 20 min served as a negative control. In one setup, latex beads were pre-treated with 1% (w/v) BSA (Sigma-Aldrich, #A9647) or 1% (w/v) β2m for 20 min at room temperature before being washed, resuspended in HBSS, and then added to the PMNs. After the 4-hour incubation, cells were washed in HBSS and run on a CytoFLEX S (Beckman Coulter). At least 10,000 events were recorded, and data was analyzed using FlowJo (v10.10.0).

Separately, 2×10^5^ PMNs were treated with pHrodo™ Red Zymosan BioParticles™ (Thermo Fisher Scientific, #P35364) either alone or with 50 µg/ml β2m or dK58β2m for 2 h at 37°C, 5% CO_2_. Zymosan particles were added at a ratio of 1:10 relative to PMNs. PMNs pre-incubated with 10 µg/ml cytochalasin D (Thermo Fisher Scientific, #PHZ1063) for 20 min were used as a control for uptake. After incubation, the cells were washed in HBSS and run on a CytoFLEX S (Beckman Coulter). At least 10,000 events were recorded, and data was analyzed using FlowJo (v10.10.0).

### Phagocytosis assay with bacteria

AP1 and KR792 strains were diluted to approximately 2×10^5^ CFU/ml, and 50 µl of the bacterial suspensions were transferred to a 96-well V-bottom plate. The bacteria were briefly centrifuged at 5000 rcf and the supernatant was discarded. Bacterial pellet was resuspended in 0.05 M pHrodo Deep Red dye (Thermo Fisher Scientific, #P35359) diluted in 0.1 M sodium bicarbonate (Thermo Fisher Scientific, #25080-060), and stained with pHrodo for 1 h in the dark at RT. After staining, bacteria were centrifuged, washed with PBS, and then resuspended in 50 µl GVB^++^ buffer (5 mM veronal buffer [pH 7.3], 140 mM NaCl, 0.1% gelatin, 1 mM MgCl_2_, and 5 mM CaCl_2_). Then, 50 µg/ml of β2m or dK58β2m, or 5 µg/ml of cytochalasin D (Sigma-Aldrich, #C8273) acting here as a negative control were added. Next, NHS was added at concentrations of 10% (for AP1) and 2% (for KR792), supplemented with 25 µg/ml of the complement inhibitor OmCI to prevent complement-mediated lysis of Gram-negative bacteria^49^. Afterwards, bacteria were mixed with PMNs at the multiplicity of infection (MOI) of 0.1. Samples were incubated at 37°C, 5% CO_2_ for 30, 60, and 120 min, respectively. After incubation, samples were placed on ice to stop the phagocytosis, centrifuged briefly, and extracellular bacteria were removed by washing PMNs with PBS. Finally, cells were resuspended in PBS and analyzed using a CytoFLEX multicolor flow cytometer (Beckman Coulter). PMNs were first gated based on the forward and sideward scatters, followed by the quantification of the pHrodo Deep Red signal of the phagocytized bacteria. At least 20,000 pHrodo-positive events were recorded.

### Flow cytometric analysis of efferocytosis

Efferocytosis of apoptotic Jurkat cells by PMNs was assessed using flow cytometry. To enable detection, PMNs and apoptotic Jurkat cells were labeled with CellTrace™ Far Red (Thermo Fisher Scientific, #C34564) and CellTrace™ Violet (Thermo Fisher Scientific, #C34557), respectively. These dyes covalently bind to free amines on and within the cells, making them fluorescently visible. Both cell types were stained with their respective dyes for 20 min at room temperature, and the staining reaction was quenched by adding 50% FBS (Corning, #35–079-CV) in RPMI.

The far red-labeled PMNs were then incubated with violet-stained apoptotic Jurkat cells, either alone or in the presence of 50 µg/ml β2m or dK58β2m, for 2 h at 37°C, 5% CO_2_. In an additional experimental setup, 10% or 25% human AB serum (Sigma-Aldrich, #H6914) was added to the PMN-Jurkat co-cultures, with or without 50 µg/ml β2m or dK58β2m. PMNs treated with 100 ng/ml recombinant human Granulocyte–Macrophage Colony-Stimulating Factor (GM-CSF) (R&D systems, #215-GM) served as a positive control. As a control for phagocytic uptake, PMNs were pre-treated with 10 µg/ml cytochalasin D (Thermo Fisher Scientific, #PHZ1063) for 20 min. After the two-hour incubation, the cells were washed in HBSS, and granulocytes were stained with anti-CD66b antibodies (R&D systems, #FAB4246G) for 30 min at 4°C in the dark. After a final wash, samples were run on a CytoFLEX S (Beckman Coulter) and at least 10,000 events were recorded. Data was analyzed using FlowJo (v10.10.0).

### Statistics

Statistical analysis was performed using GraphPad Prism 10. Normality tests (Shapiro-Wilk test) were run to determine the appropriate statistical test. Comparisons were made using repeated-measures ANOVA, mixed-effects analysis, repeated-measure two-way ANOVA, or Friedman test, followed by Dunnett’s or Tukey’s multiple comparisons test. A p-value <0.05 was considered statistically significant.

## Results

### Soluble β2m stimulates phagocytosis of carboxylate-modified latex beads by PMNs

Although β2m has been identified within the granules of neutrophils^5–7^, its exact role in this context remains unknown. We hypothesize that, like other granule-derived proteins^42^, β2m may influence neutrophil activity through autocrine and/or paracrine signaling following degranulation. Here, we studied the potential involvement of β2m in modulating neutrophil phagocytosis.

Latex beads are frequently used to study neutrophil phagocytosis without eliciting a pathogen-specific response^50,51^. We treated freshly isolated PMNs with carboxylate-modified latex beads, either alone or in combination with 50 µg/ml β2m or dK58β2m. After 4 hours, we measured the percentage of PMNs positive for the bead dye by flow cytometry. We found that β2m significantly increased phagocytosis compared to the control (from 23±8% to 31±8%), whereas dK58β2m had no significant effect (Figure 1A). Treatment with cytochalasin D, which inhibits phagocytosis, resulted in about 9±6% of PMNs staining positive for bead dye, suggesting that some beads adhered to the cell surface without being engulfed (Figure 1A).

**Figure 1:**
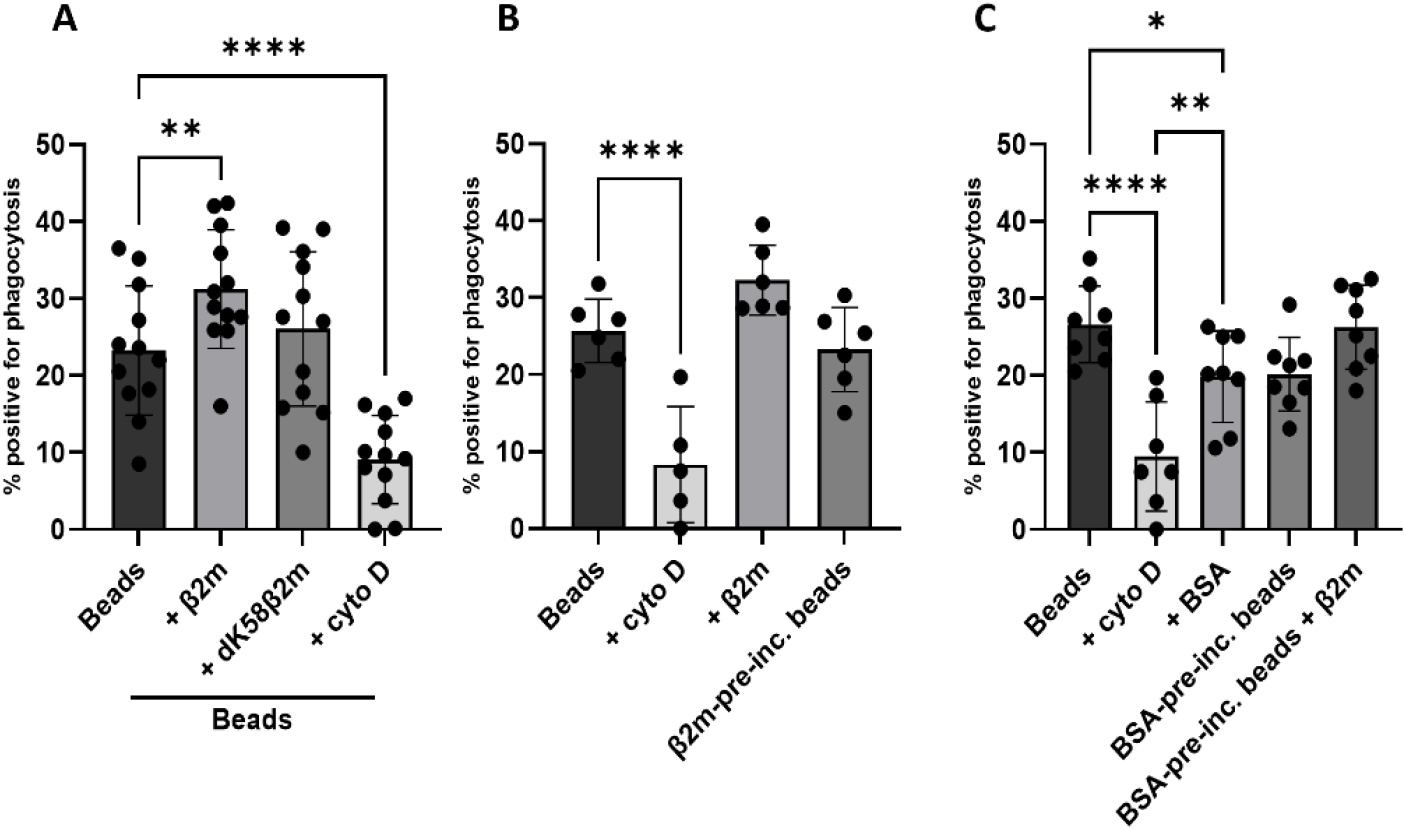
Soluble β2m enhances phagocytosis of carboxylate-modified latex beads. (A) PMNs were incubated with 50 µg/ml β2m or dK58β2m in combination with red carboxylate-modified latex beads. (B) PMNs were treated with 50 µg/ml β2m with carboxylate-modified latex beads, or with beads pre-incubated with β2m (β2m-pre-inc.) for 20 min. (C) PMNs were treated with 1% (w/v) BSA in combination with carboxylate-modified beads, or with beads pre-incubated with BSA for 20 min (BSA-pre-inc.), with or without 50 µg/ml β2m. A 10:1 bead-to-PMN ratio was used in all experiments. Pre-incubation with 10 µg/ml cytochalasin D (cyto D) served as a negative control. Results are presented as mean ± SD, as a percentage of cells positive for phagocytosis. n=12 (A), n=6 (B) and n=7 or 8 donors (C). Statistical comparisons were performed using repeated-measures ANOVA and Dunnett’s post hoc test (A), mixed-effects analysis and Dunnett’s post hoc test (B) or mixed-effects analysis and Tukey’s multiple comparisons test, with significance levels indicated as *P < 0.05, **P < 0.01, and ****P< 0.0001.

The observed increase in the percentage of PMNs positive for bead dye could have several explanations. One explanation could be that β2m acts as an opsonin, coating the beads and thereby promoting more efficient phagocytosis. To test this, we pre-incubated the beads with 1% (w/v) β2m for 20 minutes to ‘coat’ them before washing and adding them to the PMNs for 4 hours, following the same protocol used for untreated beads. We found that beads pre-incubated with β2m did not induce the same increase in phagocytosis as seen when PMNs were co-stimulated with beads and soluble β2m (Figure 1B). Instead, the uptake of β2m-coated beads was comparable to that of untreated beads. This indicates that the β2m-mediated increase in phagocytosis of latex beads is dependent on the presence of soluble β2m.

To further investigate this, we incubated the beads with BSA, a protein commonly used to reduce nonspecific interactions. This pre-incubation confirmed and supported our previous finding that β2m does not bind to the beads prior to uptake. Notably, BSA pre-treatment reduced phagocytosis compared to untreated beads (from 27±5% to 20±6%) (Figure 1C). Similarly, incubation of PMNs with carboxylate-modified beads in the presence of soluble BSA resulted in a reduced percentage of bead-positive PMNs (to 20±5%). However, when PMNs were stimulated with BSA-pre-incubated beads in the presence of soluble β2m, we still found a stimulatory effect compared to BSA-pre-incubated beads alone (from 20±6% to 26±5%) (Figure 1C).

Together, these findings suggest that β2m does not enhance phagocytosis by opsonizing latex beads. Instead, soluble β2m appears to modulate the phagocytic activity directly through its effects on PMNs.

### Both β2m and dK58β2m enhance phagocytosis of bacteria by PMNs

While latex beads serve as a simple model for studying PMN phagocytosis, they are not a physiological target. Therefore, we next examined the effects of β2m and dK58β2m on phagocytosis of two bacterial strains: the Gram-positive *S. pyogenes* AP1 and the Gram-negative *A. baumannii* clinical isolate KR792. Both strains were labeled with pHrodo deep red dye to monitor uptake via flow cytometry.

Freshly isolated PMNs were incubated with each bacterial strain at an MOI of 0.1, either alone or in the presence of 50 µg/ml of β2m or dK58β2m. Both β2m and dK58β2m significantly enhanced the uptake of AP1 and KR792 compared to untreated controls (Figure 2A and B). This stimulatory effect was time-dependent, with no significant change detected after 30 minutes, but a clear increase after 60 minutes and 120 minutes for both proteins. Importantly, this enhancement was consistent for both bacterial strains. For AP1, β2m induced a maximum increase in uptake of 4.4±0.6-fold, while dK58β2m led to a 4.8±0.6-fold increase after 2 hours. For KR792, peak uptake occurred at 60 minutes, with β2m and dK58β2m increasing phagocytosis by 3.6±1.0-fold and 2.9±0.7-fold, respectively. Notably, after 2 hours of incubation with KR792, β2m treatment resulted in significantly higher uptake compared to dK58β2m (p=0.004). In summary, β2m and dK58β2m promote bacterial phagocytosis by PMNs in a time-dependent manner.

**Figure 2:**
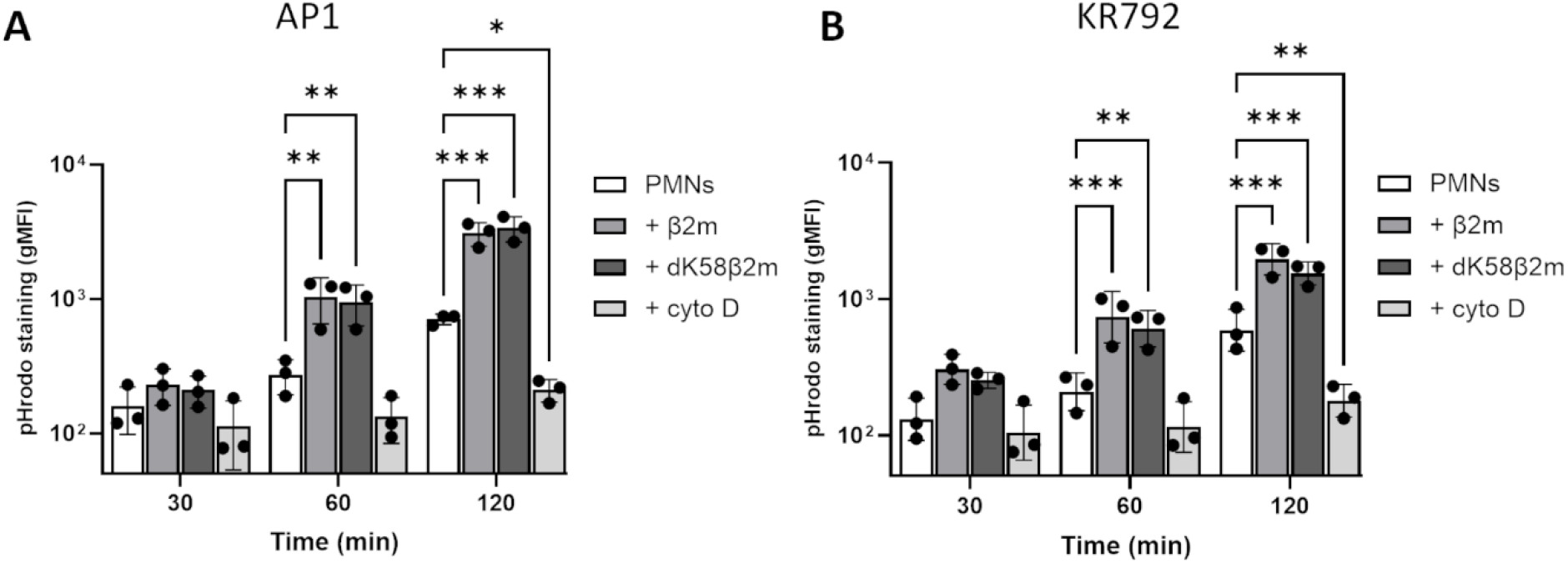
β2m and dK58β2m increase phagocytosis of *Streptococcus pyogenes* strain AP1 and *Acinetobacter baumannii* clinical isolate KR792 by PMNs. PMNs were treated with pHrodo-labelled bacteria (AP1 or KR792) at a MOI of 0.1, 10% (for AP1) or 2% (for KR792) NHS, 25 µg/ml of the complement inhibitor OmCI (to prevent lysis of Gram-negative KR792), with or without 50 µg/ml β2m or dK58β2m for 30, 60, or 120 min. Treatment with 5 µg/ml cytochalasin D (cyto D) was used as a negative control for phagocytosis. Results are expressed as mean ± SD, as gMFI (n=3 donors). Statistical significance is indicated by *P < 0.05, **P < 0.01, and ***P < 0.001. Group comparisons were performed using repeated-measure two-way ANOVA and Dunnett’s post hoc test.

We also examined phagocytosis of another target, pHrodo-labelled zymosan bioparticles. In this experimental setup, we did not find an effect on uptake with the two β2m forms (Supplementary figure 2).

### β2m and dK58β2m also stimulate efferocytosis of apoptotic Jurkat cells by CD66b-positive granulocytes

In addition to engulfing pathogens, phagocytic cells like neutrophils can clear apoptotic cells through efferocytosis^38,39,41^. Unlike phagocytosis, efferocytosis primarily supports tissue homeostasis rather than contributing to inflammation. To explore whether β2m and dK58β2m influence efferocytosis, we induced apoptosis in Jurkat cells by treating them with 0.8 µM staurosporine for 24 hours. These apoptotic cells were then labeled with CellTrace™ Violet to track their uptake by CD66b-positive granulocytes, which were stained with CellTrace™ Far Red. After co-culturing for two hours, we found that both β2m and dK58β2m increased efferocytosis in a dose-dependent manner, peaking at 50 µg/ml (Figure 3A). The percentage of cells double-positive for both CellTrace dyes was found to be similar to treatment with 100 ng/ml GM-CSF, a known inducer of phagocytosis and efferocytosis in neutrophils^40,52^. When testing the combined effect of β2m or dK58β2m with GM-CSF, we found no further enhancement beyond the effect of each stimulus alone (Figure 3B).

**Figure 3:**
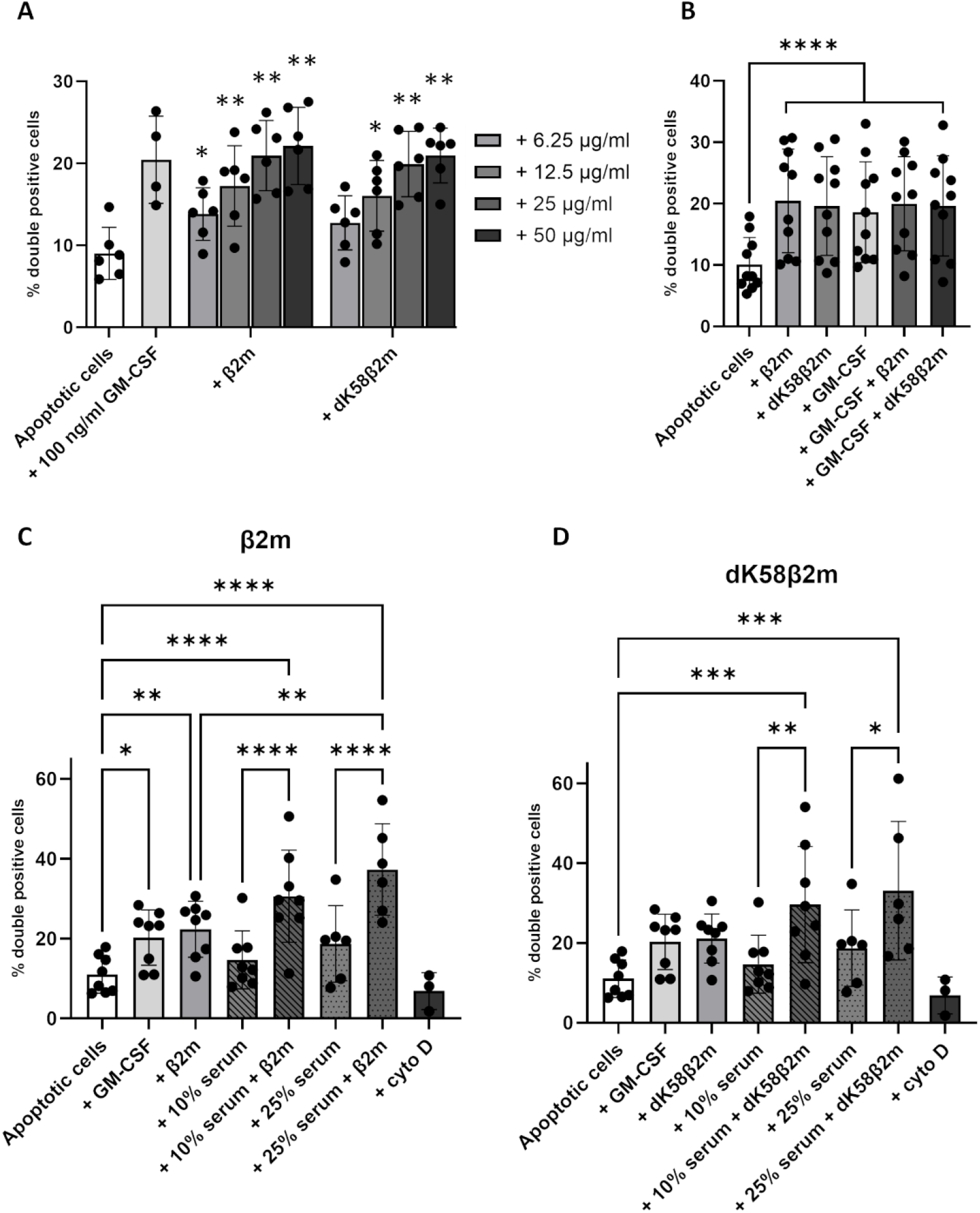
β2m and dK58β2m both enhance efferocytosis of apoptotic Jurkat cells by CD66b-positive granulocytes. Efferocytosis of apoptotic Jurkat cells labeled with CellTrace™ Violet by CD66b-positive granulocytes stained with CellTrace™ Far Red was examined under various conditions. (A) PMNs were exposed to varying concentrations of β2m or dK58β2m in the presence of apoptotic Jurkat cells. (B) PMNs were treated with apoptotic cells alone or in combination with 50 µg/ml β2m, 50 µg/ml dK58β2m, 100 ng/ml GM-CSF, or combinations thereof. (C-D) PMNs were treated with apoptotic Jurkat cells with 50 µg/ml β2m (C) or dK58β2m (D) combined with 10 or 25% human AB serum. In all experiments, a 4:1 apoptotic cell-to-PMN ratio was used. Treatment with 100 ng/ml GM-CSF served as a positive control, and pre-incubation with 10 µg/ml cytochalasin D (cyto D) served as a negative control. Results are shown as mean ± SD of double-positive PMNs. Sample sizes were n=6 or n=4 (A), n=10 (B), and n=8 or 3 donors (C and D). Group comparisons were analyzed using mixed-effects analysis and Tukey’s multiple comparisons test (A, C, and D) or repeated-measures ANOVA and Tukey’s multiplecomparisons test(B). Statistical significance is indicated by *P < 0.05 and **P < 0.01, ***P < 0.001, ****P < 0.0001.

Since serum is also recognized to stimulate efferocytosis^40^, we performed experiments with 10% and 25% human AB serum. Co-stimulation of PMNs with β2m and human serum significantly increased the percentage of double-positive cells compared to serum alone (Figure 3C). Although this combination did not significantly exceed β2m treatment alone, there was a trend toward enhanced efferocytosis when β2m and serum were combined. For dK58β2m, only co-stimulation with 10% human serum resulted in significantly higher efferocytosis compared to serum alone (Figure 3D).

## Discussion

In the present study, we show that both β2m and its cleaved variant dK58β2m not only promote phagocytosis of *S. pyogenes* AP1 and *A. baumannii* KR792 but also stimulate efferocytosis of apoptotic cells. Interestingly, only β2m, but not dK58β2m, increases the uptake of latex beads, and this effect depends on the presence of soluble β2m.

Phagocytosis of foreign particles is a key function of the innate immune system, critical for controlling inflammation and infection^53^. Our experiments with latex beads revealed that soluble β2m stimulates phagocytosis of these particles, whereas dK58β2m did not. To determine if β2m enhanced uptake by directly binding to the beads, we pre-incubated beads with β2m, but this had no effect on phagocytosis. When beads were coated with BSA to reduce nonspecific binding, their uptake decreased, consistent with previous reports^54^. Interestingly, adding soluble β2m alongside BSA-coated beads also showed a stimulatory effect of β2m. This suggests that the stimulatory effect of β2m requires its soluble form and is not due to β2m binding to or opsonizing the beads for a more efficient phagocytosis.

We also investigated phagocytosis of two bacterial strains, *S. pyogenes* AP1 and *A. baumannii* KR792, and found that both β2m and dK58β2m enhance this process. Since *S. pyogenes* is a Gram-positive bacterial strain and *A. baumannii* is Gram-negative, these findings suggest that the stimulatory effect of β2m is independent of the bacterial Gram status. To follow bacterial phagocytosis, we labelled the bacteria with pHrodo, a pH-sensitive dye whose fluorescence increases in acidic environments such as that of the phagolysosome. Therefore, an increase in fluorescence intensity (measured as gMFI) reflects either a higher number of bacteria engulfed by PMNs, more efficient delivery of bacteria to the phagolysosome, or a combination of both.

Interestingly, β2m knockout (KO) mice exhibit impaired bacterial clearance and increased mortality following intravenous infection with the Gram-negative bacterial strain *Klebsiella pneumoniae (K. pneumoniae)*^55^. This effect is unlikely to result from the role of β2m in MHC class I expression, as TAP-1 KO mice, which like β2m KO mice lack MHC class I and CD8^+^T cells, showed mortality rates similar to wild-type (WT) mice. This interpretation is further supported by the observation that most deaths occurred early, between day 2 and 4 post-infection, before full activation of the adaptive immune response. In contrast, MHC class I–related functions of β2m have been shown to be important for protection against *Mycobacterium tuberculosis* in mice^56^. In this study, a decrease in mortality was first observed several weeks post-infection and was shown to related to the adaptive immune response^56^.

While we did not test the effect of β2m on phagocytosis of *K. pneumoniae*, we did find that β2m can promote phagocytosis of another Gram-negative bacterium, *A. baumannii*. This raises the possibility that the higher bacterial load of *K. pneumoniae* observed in β2m KO mice could stem from reduced phagocytic function of PMNs. Supporting this, PMNs isolated from β2m KO mice have been shown to exhibit impaired phagocytosis of IgG1-opsonized Gram-positive *Streptococcus pneumoniae (S. pneumoniae)*^57^. The decreased uptake was linked to the absence of functional neonatal Fc receptors (FcRn), which require β2m for stability, as PMNs from FcRn α-chain KO mice showed a similar impairment. However, our findings raise the possibility that β2m itself may have a stimulatory role in phagocytosis of *S. pneumoniae* independent of FcRn.

In addition to their well-established role in engulfing pathogens, neutrophils are increasingly recognized for their involvement in clearing apoptotic cells, a process essential for maintaining tissue homeostasis and regulating inflammation^40,41^. In this study, we show that both β2m and dK58β2m can enhance efferocytosis of apoptotic Jurkat cells by PMNs in a dose-dependent manner. Notably, 50 µg/ml of β2m and dK58β2m induced efferocytosis at levels comparable to treatment with 100 ng/ml of GM-CSF, a known stimulator of efferocytosis^40^. Interestingly, co-stimulation with both β2m and GM-CSF did not produce an additive effect, suggesting that these molecules may promote efferocytosis via a shared mechanism or signaling pathway. In contrast, co-stimulation with human serum showed a tendency toward an additive effect when combined with either β2m or dK58β2m. GM-CSF is known to upregulate bridging molecules such as MFG-E8, which binds phosphatidylserine (an “eat-me” signal) on apoptotic cells and interacts with integrins on phagocytes such as macrophages and dendritic cells^58,59^. It also stimulates expression of MerTK on these phagocytes, whose ligand GAS6 similarly binds phosphatidylserine to enhance apoptotic cell recognition and engulfment^60,61^. It would be interesting to investigate whether β2m facilitates efferocytosis through a similar mechanism in neutrophils.

Since both β2m and dK58β2m augment phagocytosis of the two bacterial strains and efferocytosis of apoptotic cells, the stimulatory activity of dK58β2m likely reflects conserved functions of other parts of β2m rather than novel properties gained through the proteolytic cleavage.

The concentrations of β2m used in this study are substantially higher than typical serum levels in healthy individuals, which are generally below 2 mg/L^62,63^. Elevated β2m serum levels are associated with various inflammatory diseases and malignancies, reaching up to 10 mg/L^63^, and in chronic kidney disease, levels as high as 75 mg/L have been reported^64^. In local inflamed areas, such as in crevicular fluid in patients with periodontitis, the concentration of β2m can reach 100 mg/L^65^. Our use of up to 50 mg/L β2m thus falls within the range observed in a number of pathological states. Moreover, serum concentrations may not accurately reflect β2m levels within local tissue environments. For example, β2m concentrations in synovial fluid from patients with rheumatoid arthritis often exceed serum levels^66^. Therefore, it is plausible that local β2m concentrations, especially within confined compartments such as synovial fluid during inflammation, may surpass those measured in serum.

In summary, we show that β2m and dK58β2m enhance phagocytosis of *S. pyogenes* AP1 and *A. baumannii* KR792, as well as efferocytosis of apoptotic cells by PMNs. Additionally, the native β2m, but not dK58β2m, stimulates the uptake of latex beads. Together, these results establish a role for β2m in promoting PMN-mediated clearance of both pathogens and dying cells.

## CRediT Authorship contribution statement

**Sofie Espersen Poulsen**: Conceptualization, Methodology, Formal analysis, Investigation, Writing - Original Draft, Visualization, Data Curation. **Michal Magda:** Conceptualization, Methodology, Formal analysis, Writing - Review & Editing. **Anna Blom:** Conceptualization, Methodology, Investigation, Writing - Review & Editing. **Mogens Holst Nissen:** Conceptualization, Methodology, Writing - Review & Editing, Supervision, Project administration, Funding acquisition, Data Curation.

## Acknowledgements

Thanks to Lotte Nielsen for providing the construct for human beta 2-microglobulin used in the ExpiCHO Expression System and Nethe Thane Nielsen for technical support.

