## Supplemental Figures for "Beta-2-microglobulin stimulates neutrophil phagocytosis of bacteria and apoptotic cells"

### Supporting information

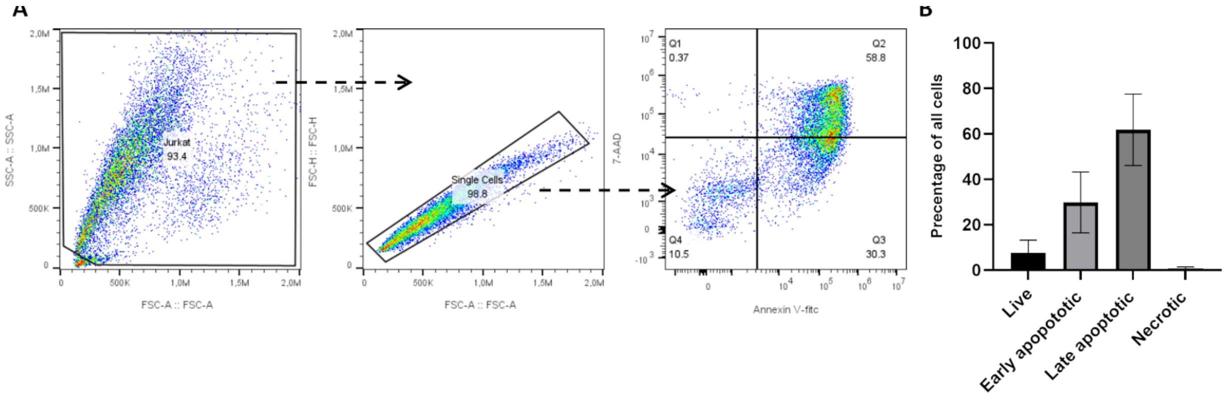


**Figure S1 – Degree of cell death in staurosporine-treated Jurkat cells.** Jurkat cells were treated with 0.8 µM staurosporine for 24 h, after which the degree of cell death was assessed. (A) Representative gating strategy for identifying apoptotic Jurkat cells stained with annexin V (FITC) and 7-AAD. (B) Results are expressed as mean ± SD, representing the percentage of cells in each cellular stage (n=8 independent repeats).

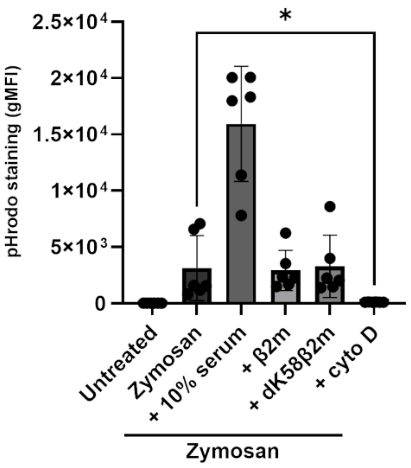


**Figure S2 - β2m and dK58β2m does not affect phagocytosis of zymosan bioparticles.** PMNs were treated with 50 µg/ml β2m or dK58β2m in combination with red pHrodo™-zymosan bioparticles at a 10:1 zymosan-to-PMN ratio. Treatment with 10% human AB serum acted as a positive control, while pre-incubation with 10 µg/ml cytochalasin D (cyto D) served as a negative control. Results are presented as mean ± SD, as geometric mean fluorescence (gMFI) (n=6 donors). Statistical analysis was conducted using the Friedman test and Dunn's multiple comparisons test, with $^*P < 0.05$.

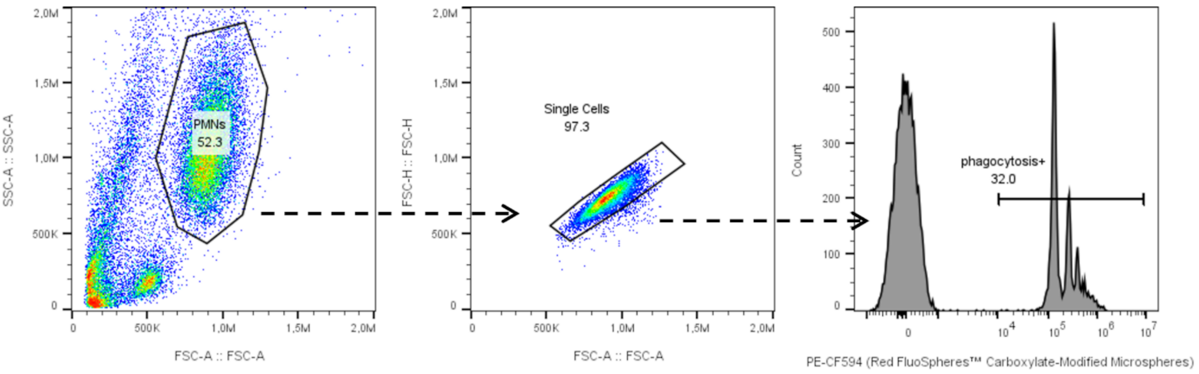


**Figure S3 – Representative gating strategy for identifying PMNs positive for phagocytosis of Red FluoSpheres$^{TM}$ Carboxylate-modified microspheres (latex beads) as shown in figure 1.**

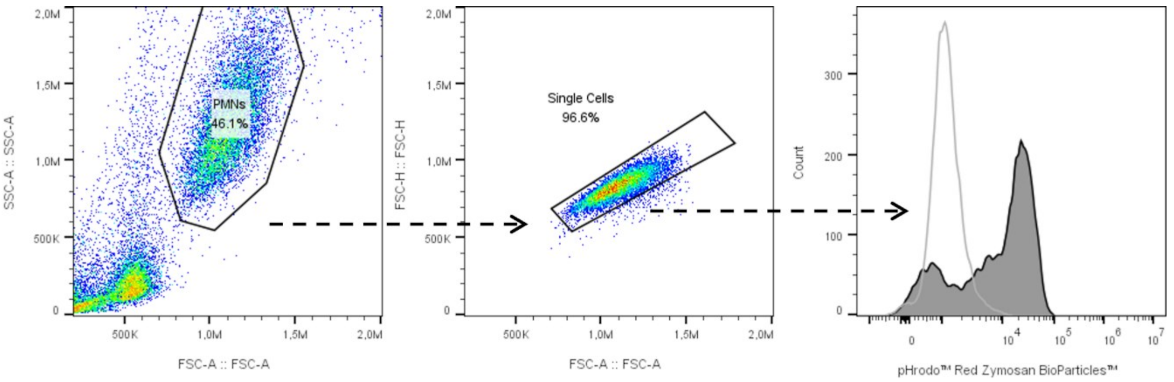


**Figure S4 – Representative gating strategy for determining gMFI for PMNs treated with pHrodo$^{TM}$ red zymosan bioparticles as illustrated in supplementary figure 1.** The dark grey filled histogram shows pHrodo-staining in PMNs treated with pHrodo$^{TM}$ red zymosan bioparticles, while the light grey line shows fluorescence of pHrodo$^{TM}$ red zymosan bioparticles alone (in HBSS).

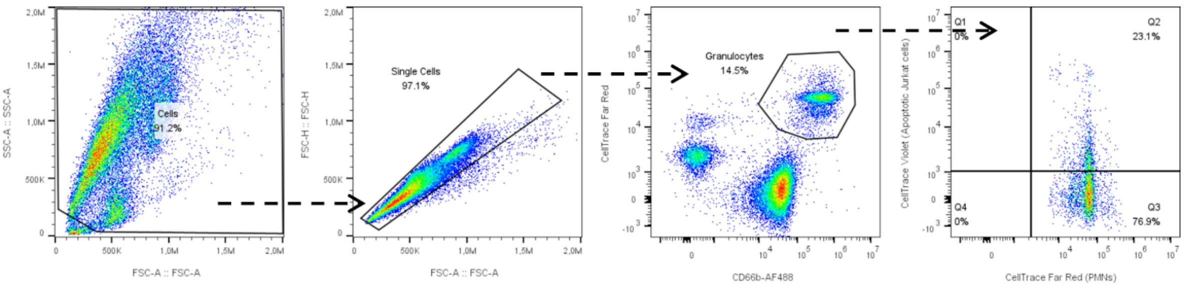


**Figure S5 – Representative gating strategy for determining the percentage of CD66-positive granulocytes double positive for CellTrace far red (e.g., PMN marker) and CellTrace violet (e.g., apoptotic Jurkat marker) as shown in figure 3.**

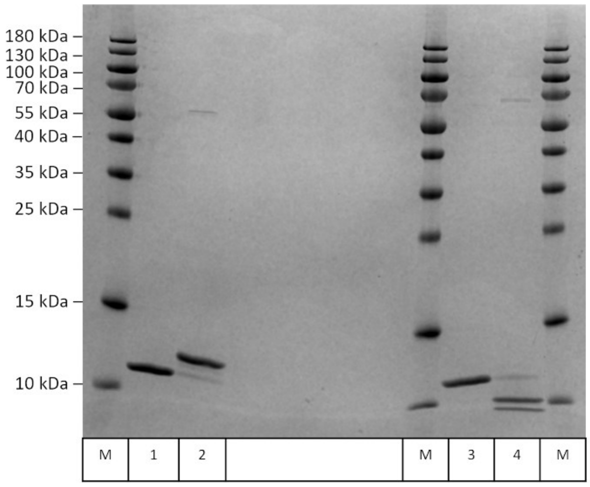


| | |
|---|---|
| M | Molecular marker |
| 1 | β2m non-reduced |
| 2 | dK58β2m non-reduced |
| 3 | β2m reduced |
| 4 | dK58β2m reduced |

**Figure S6 – Analysis of β2m and dK58β2m under non-reduced and reduced conditions.** Equal amount of purified β2m and dK58β2m were run on an SDS-PAGE and total protein levels were visualized by InstantBlue$^{TM}$ staining. Under reducing conditions, dK58β2m separates into two distinct chains, consistent with cleavage occurring within the disulfide loop of β2m. Trace amounts of native β2m can be seen present in the dK58β2m samples.
